# Persistence of Hippocampal and Striatal Multivoxel Patterns During Awake Rest after Motor Sequence Learning

**DOI:** 10.1101/2021.06.29.450290

**Authors:** Bradley R. King, Mareike A. Gann, Dante Mantini, Julien Doyon, Geneviève Albouy

## Abstract

Memory consolidation, the process by which newly encoded and fragile memories become more robust, is thought to be supported by the reactivation of brain regions - including the hippocampus - during post-learning rest. While hippocampal reactivations have been demonstrated in humans in the declarative memory domain, it remains unknown whether such a process takes place after motor learning. Using multivariate analyses of task-related and resting state fMRI data, here we show that patterns of brain activity within both the hippocampus and striatum elicited during motor learning persist into post-learning rest, indicative of reactivation of learning-related neural activity patterns. Moreover, results indicate that hippocampal reactivation reflects the spatial representation of the learned motor sequence. These results thus provide insights into the functional significance of neural reactivation after motor sequence learning.

## Introduction

Memory consolidation is thought to be supported by the replay of hippocampal activity during subsequent offline episodes, i.e. during post-encoding rest ^1–3^. Hippocampal replay, initially reported in the rodent literature, has recently been translated to human research with the development of multivariate pattern analyses of functional magnetic resonance imaging (fMRI) data ^1^. Using these approaches, several studies have shown that hippocampal brain patterns elicited by declarative learning can persist into subsequent rest episodes and that this spontaneous memory reactivation is related to better memory [e.g., ^4–6^].

Despite these recent advances, several critical knowledge gaps remain. First, hippocampal reactivations have predominantly been limited to examinations in the context of declarative memory, which is surprising given that the role of the hippocampus in motor (sequence) memory processes is now well established [see ^7,8^ for reviews]. Specifically, previous research has not only shown that the hippocampus is recruited during both implicit and explicit versions of motor sequence learning (MSL) [e.g., ^9–14^], but also that hippocampal activity and connectivity patterns elicited by task practice are linked to subsequent motor memory consolidation ^10,15^. In line with this, research using multivariate approaches has shown that task-related hippocampal patterns can discriminate newly learned as compared to consolidated motor sequences ^16^. While there is recent evidence to suggest that the hippocampus is involved in micro-offline motor memory processes [i.e., fast consolidation occurring during short rest intervals between practice blocks; see ^17,18^], persistence of hippocampal patterns after motor learning has never been reported. Second, whereas previous rodent research has demonstrated that hippocampal replays consolidate previously learned spatial relationships (e.g., ^19^), the functional significance of hippocampal reactivation in humans is poorly understood.

The current study aims to address these knowledge gaps. Resting state (RS) scans from 55 young healthy participants were acquired immediately before and after participants completed a motor sequence learning (MSL) task inside the MR scanner (Task A; Figure 1A). Participants were then subsequently assigned to one of two experimental groups and completed a MSL task variant (Task B) that probed the allocentric (i.e., spatial) or egocentric (i.e., motor) representation of the acquired motor sequence that are known to be dependent on hippocampo- or striato-cortical regions, respectively ^20^. Task B was then followed by a third RS scan. In three regions of interest (ROIs) known to be involved in motor memory consolidation (the hippocampus, striatum and primary motor cortex (M1)), we examined: a) whether multivoxel patterns of neural activity showed evidence of persistence into post-learning rest; and, b) the functional significance of pattern persistence (i.e. whether pattern persistence reflected specific representations of the acquired motor sequence). Based on the aforementioned literature on the hippocampus as well as previous evidence demonstrating that (i) the hippocampus can lead striatal replay ^21^ and (ii) task-related patterns in the primary motor cortex (M1) can be subsequently reactivated offline ^22,23^, the following hypotheses were made. We predicted that response patterns elicited by motor task practice in our three ROIs would persist into post-learning rest, reflecting reactivation of learning-related patterns. Moreover, and in line with our earlier research^20^, we postulated that pattern persistence in the hippocampus would reflect the reactivation of the spatial features of the task whereas persistence in the striatum and M1 would reflect the reinstatement of the motoric component of the task.

**Figure 1:**
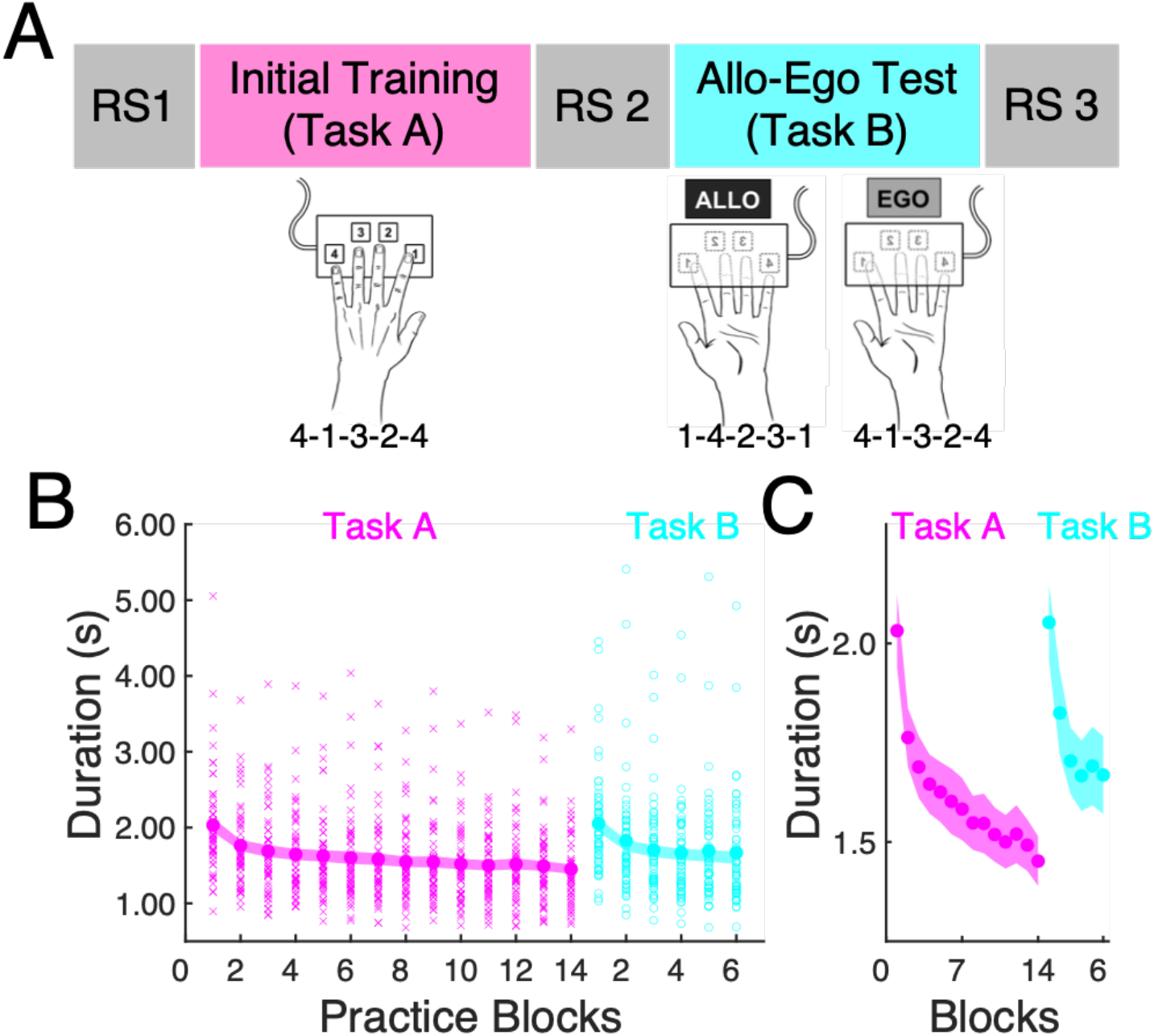
Experimental Design and Behavioral Results. **A.** Resting state (RS) scans were acquired immediately prior to (RS1) and after (RS2) the initial training of a motor sequence learning task (Task A) in 55 young healthy individuals as well as following (RS3) a test session (Task B) assessing allocentric (Allo; n = 26) or egocentric (Ego; n = 29) sequence representations. Both tasks consisted of explicitly known 5-element sequences completed with the left non-dominant hand (1 = index finger; 2 = middle finger, 3 = ring finger, 4 = little finger). The egocentric representation Task B was assessed by having the participants complete the same sequence of finger presses as the initial training (i.e., 4-1-3-2-4); however, the spatial locations of the movement responses were now novel as both the keyboard and hand were inverted. For the allocentric representation, the spatial representation of the sequence was preserved by having the participants complete the mirrored finger sequence (i.e., 1-4-2-3-1) after inversion of the keyboard and hand. Note that all tasks and RS runs were completed inside the MRI scanner. **B.** Participants exhibited significant reductions in the duration to complete a correct sequence (in s) across practice blocks of Task A (magenta), reflective of learning the sequence of movements. Similar results were obtained for Task B (cyan) that followed RS2. As performance speed did not differ between allocentric and egocentric representations of the motor sequence task (see main text for statistical details), the plots above collapsed across the two experimental groups. Large circles represent means, shaded regions represent SEM, small x’s and o’s denote individual data for training and test runs, respectively. **C**. Zoomed-in version of behavioral data depicted in panel B to better represent reductions in movement speed as a function of practice.

## Results

### Behavior

Participants performed an explicitly known five-element finger sequence with their non-dominant (left) hand while positioned supine in the fMRI scanner and brain images were recorded (Figure 1A). During the initial training session (Task A), the time to perform a correct sequence (in seconds) decreased across blocks of practice (F_(13,689)_ = 36.8, Greenhouse-Geisser corrected p-value < 0.001, partial η^2^ = 0.404; Figure 1B/1C). Following the post-training RS scan, participants were divided into two groups and completed 6 additional blocks of a task variant (Task B) that probed either the allocentric (i.e., spatial) or egocentric (i.e., motor) representation of the acquired motor sequence (see ^20^ and methods below). Similar to Task A, the time to complete a sequence for Task B significantly decreased as a function of practice blocks (F_(5,270)_ = 40.4, corrected p-value < 0.001, partial η^2^ = 0.428). While the brain responses subtending these different task representations are known to be distinct ^20^, our analyses of the current sample was consistent with previous research ^20,24^ demonstrating no differences in movement speed between the two task variants (group main effect: F(i,53) = 2.56, corrected p-value = 0.12, partial η^2^ = 0.046; group x block interaction: F_(5,265)_ = 0.32, corrected p-value = 0.86, partial η^2^ = 0.006). These results collectively demonstrate the expected finding that participants exhibited evidence of motor sequence learning in both tasks, as reflected by improvements in movement speed.

### Persistence of task patterns following Task A

We first examined whether brain patterns during Task A (i.e., initial sequence training) within our three ROIs (Figure 2B) showed evidence of persistence into post-learning rest. To do so, we computed for each ROI and for each fMRI run of interest (i.e., RS1, Task A and RS2) a multivoxel correlation structure (MVCS; see Figure 2A) reflecting the *multivoxel activity pattern* of the ROI ^4^. Next, for each ROI, we measured the similarity between rest- and task-related multivoxel patterns, defined as the r-to-z transformed correlation between the two MVCS matrices. We then tested whether RS1/Task A and Task A/RS2 similarity indices differed, with greater Task A/RS2 similarity indicative of persistence of the task-related pattern into subsequent rest (i.e., similarity with RS1 serves as a within-subject control) ^4^. Results showed that the multivoxel response patterns within the hippocampus (HC) and putamen (Put) during Task A were significantly more similar to activity patterns in the post-training RS2 scan as compared to the pre-training RS1 scan (HC: (t(54) = 2.86, p(unc) = 0.006, p(FDR) = 0.018, Cohen’s d = 0.39; Put: t(54) = 2.45, p(unc) = 0.018, p(FDR) = 0.027, Cohen’s d = 0.33; Figure 2C). In contrast, the similarity of M1 activity patterns between Task A and the RS scans did not differ between pre- and post-training RS (t(54) = 1.37, p(unc) = 0.18; Cohen’s d = 0.18; Figure 2C). These results indicate that task-related patterns within the hippocampus and striatum – but not in M1– significantly persisted into rest after initial motor sequence training.

**Figure 2:**
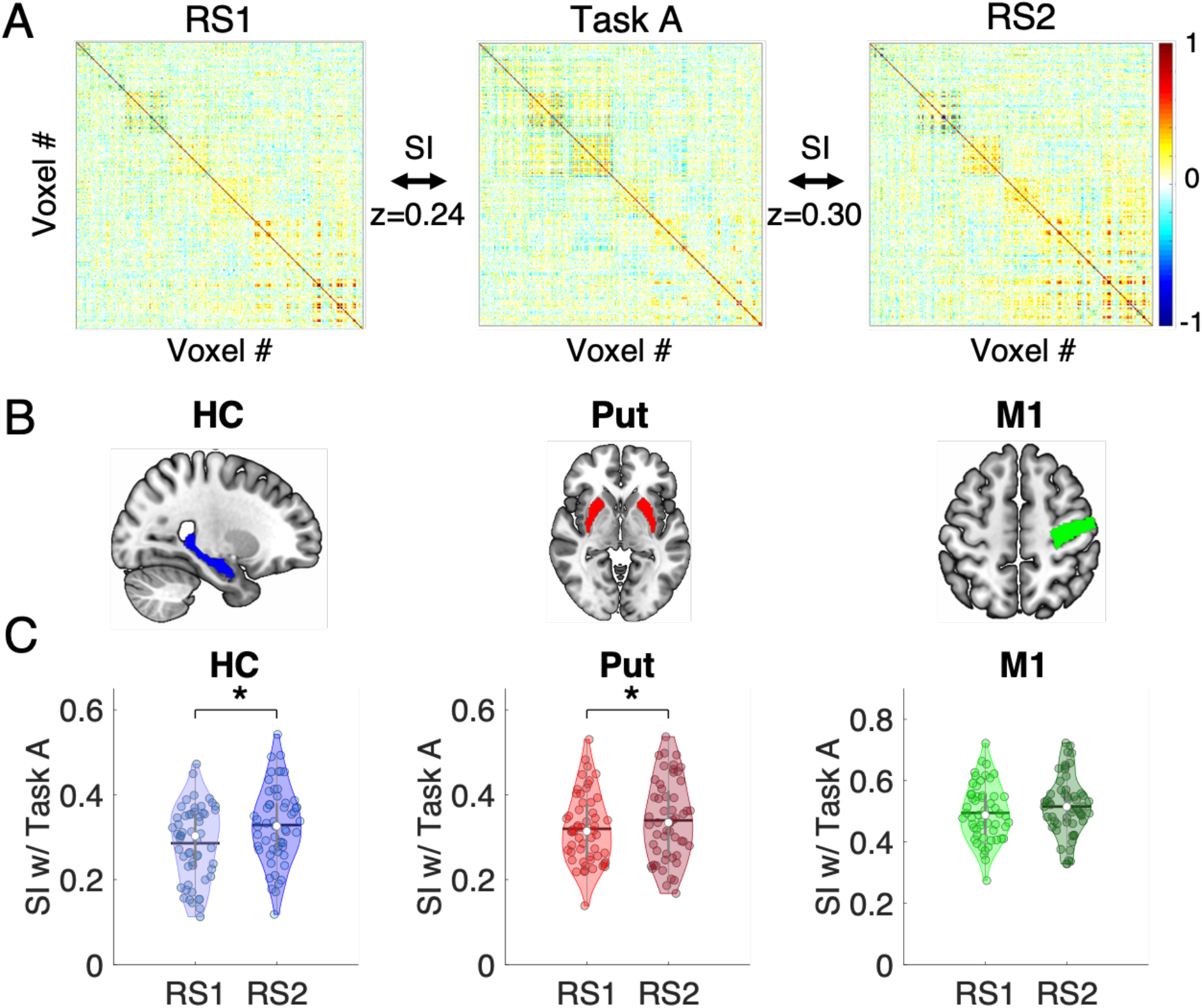
Pattern persistence following initial learning. (**A**) MVCS matrices are depicted for an exemplar ROI and participant. Each matrix shows the correlation between each of the n voxels of the ROI with all the other voxels of the ROI during the pre-training resting state (RS1), Task A (i.e., motor sequence learning (MSL) training) and post-training resting state (RS2) runs. The similarity index (SI) between two matrices (RS1 vs. Task A and Task A vs. RS2) is defined as the r-to-z transformed correlation between each pair of MVCS matrices. RS1/Task A and Task A/RS2 SIs were compared to assess whether task-related patterns persisted significantly into post-learning rest (results presented in panel C). (**B**) Depiction of the hippocampus (HC), putamen (Put) and primary motor cortex (M1) ROIs overlaid on the MNI 152 template as part of the software MRIcroGL available at https://www.mccauslandcenter.sc.edu/mricrosl. (**C**) Violin plots ^25^ depicting the Similarity Indices (SI; Fisher Z-transformed correlation coefficients) between RS1 and Task A as well as between Task A and RS2 for the HC, Put and M1. For the HC and Put, the pattern of neural responses observed during Task A persisted into the subsequent rest, reflective of reactivation of learning-related neural activity. For all violin plots, individual data points are shown as small colored circles jittered on the horizontal axis within the respective plot to increase visualization (N = 55); white circles and horizontal black lines depict group medians and means, respectively. * indicates p < 0.05 after FDR correction for multiple comparisons.

### Functional significance of pattern persistence following Task A

Task B was designed to probe the different task representations (i.e., allocentric and egocentric) that are known to develop during initial motor sequence learning (i.e., during Task A practice, see ^20,26^). Here, we assessed whether post-learning RS2 patterns reflected allocentric and/or egocentric representations of the task. To do so, we tested whether RS2/Task B similarity indices differed between allocentric and egocentric task conditions in the two ROIs showing pattern persistence (i.e., the hippocampus and putamen). Results for the hippocampal ROI showed that the similarity between RS2 and Task B was significantly greater for the allocentric as compared to the egocentric representation (t(52) =2.67, p(unc) = 0.010, p(FDR) = 0.020, Cohen’s d = 0.73; Figure 3). This suggests that the hippocampal multivoxel patterns reactivated during post-training RS2 reflect the allocentric representation of the sequence task. For the putamen, the difference between the allocentric and egocentric groups did not statistically differ (t(52) = 1.74, p(unc) = 0.09, Cohen’s d = 0.47), suggesting that the persistence of learning-related activity in the putamen does not differentially reflect allocentric or egocentric representations of the motor sequence.

**Figure 3:**
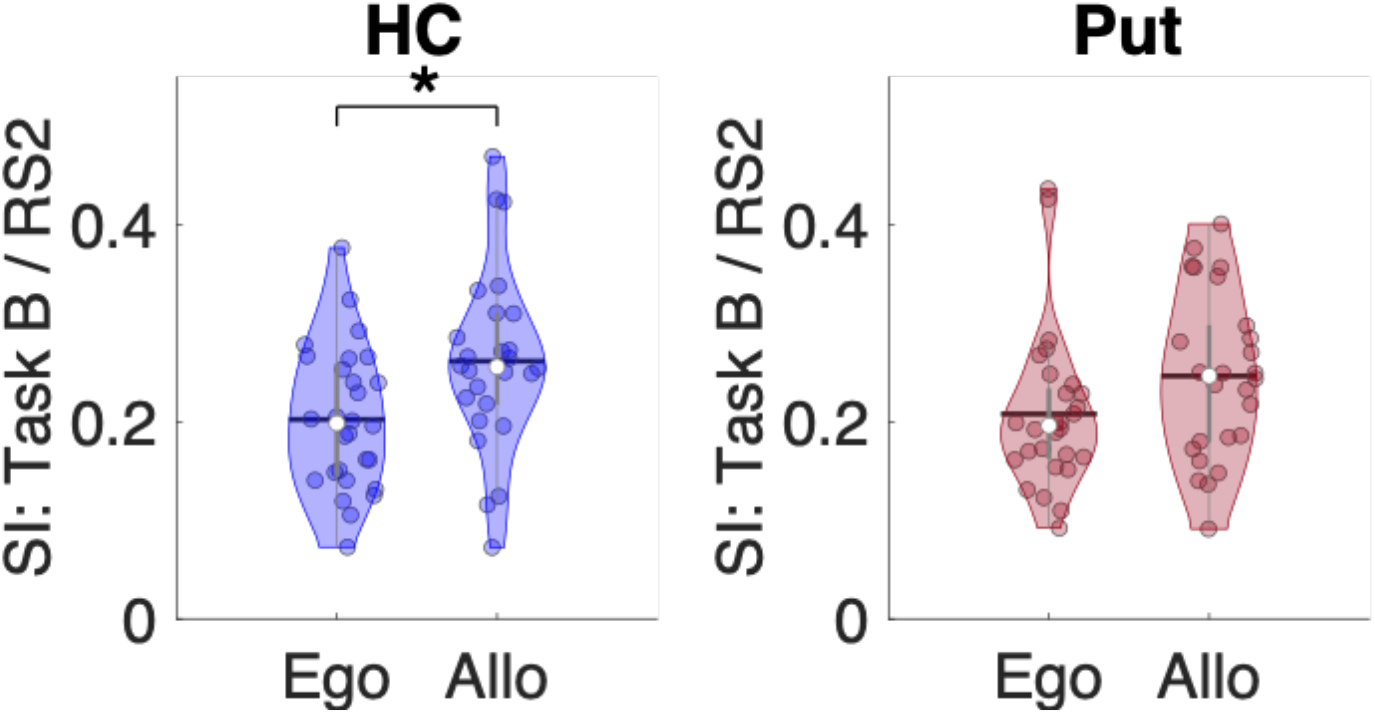
Functional significance of pattern persistence following Task A. Violin plots ^25^ depicting the Similarity Indices (SI) between MVCS matrices from post-training RS (i.e., RS2) and Task B for the hippocampus (HC; blue) and putamen (Put; red) separately for the egocentric (Ego) and allocentric (Allo) groups. For the HC, SI was greater for the allocentric than the egocentric group, suggesting pattern persistence during RS2 (following initial learning) reflects the reactivation of the allocentric representation of the motor sequence. There were no group differences for the Put. For all violin plots, individual data points are shown as small colored circles jittered on the horizontal axis within the respective plot to increase visualization (N = 28 and 26 in Ego and Allo groups, respectively; one subject in Ego was excluded due to missing data); white circles and horizontal black lines depict group medians and means, respectively. * indicates p < 0.05 after FDR correction for multiple comparisons.

### Persistence of task patterns following Task B

Our next analyses examined whether multivoxel patterns elicited by Task B showed evidence of persistence into the subsequent rest interval. To do so, we tested whether RS2/Task B and Task B/RS3 similarity indices differed, with greater Task B/RS3 similarity indicative of persistence of the task-related pattern into subsequent rest. Results demonstrated that hippocampal multivoxel patterns elicited by Task B (irrespective of allocentric/egocentric representations) indeed persisted into the RS3 epoch. Specifically, the Task B pattern was more similar to RS3 than RS2 (F(1,53) = 5.44, p(unc) = 0.024, p(FDR) = 0.048, Cohen’s d = 0.32; Figure 4). For the putamen, the multivoxel pattern during the Task B did not persist into the subsequent rest interval (F(1,53) = 0.22, p(unc) = 0.64, Cohen’s d = 0.06).

**Figure 4:**
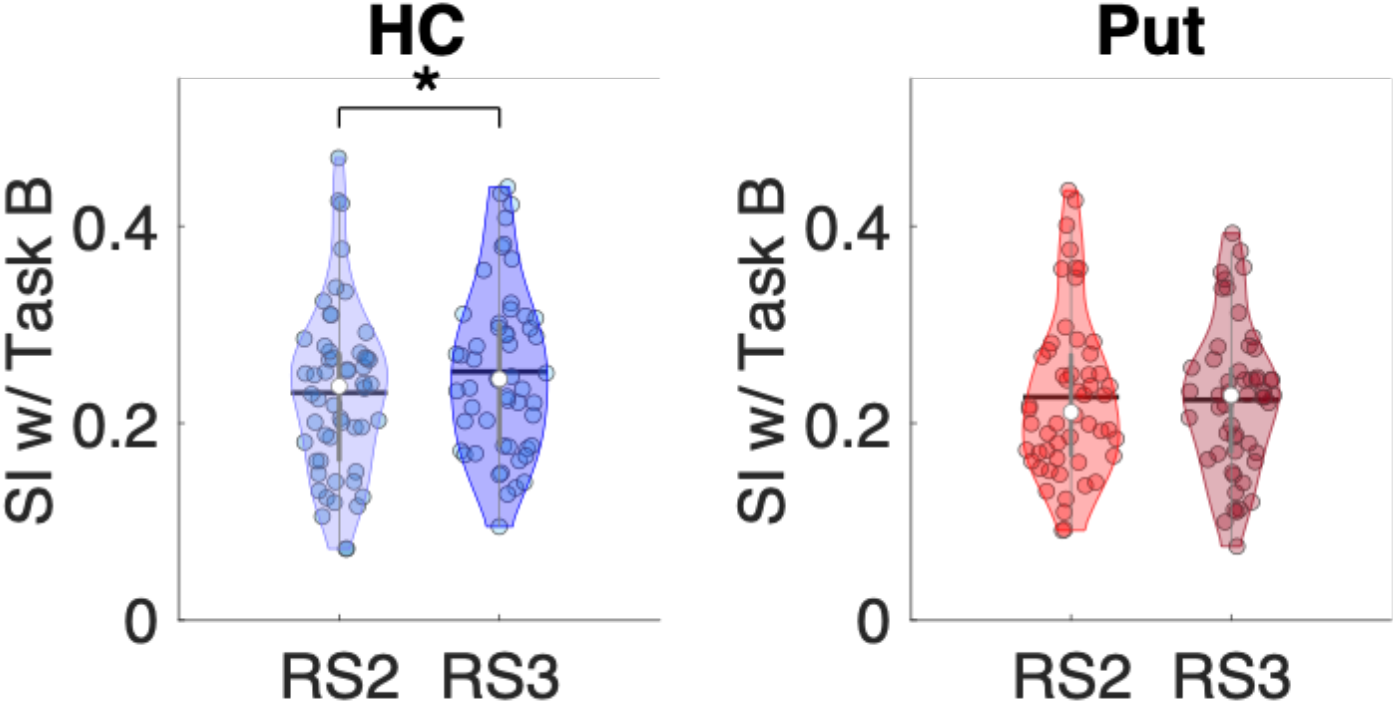
Persistence of task patterns following Task B. Violin plots ^25^ depicting the Similarity Indices (SI) between MVCS matrices from the Task B and those during RS2 and RS3 for the hippocampus (HC; blue) and putamen (Put; red). For the HC, the pattern of neural responses during Task B was more similar to RS3 than RS2, indicative of persistence. The patterns of neural responses in the Put during Task B were equally similar to the patterns observed during the preceding rest (RS2) and the subsequent rest (RS3) for both allocentric and egocentric conditions. For all violin plots, individual data points are shown as small colored circles jittered on the horizontal axis within the respective plot to increase visualization (N = 54; one subject was excluded due to missing data); white circles and horizontal black lines depict group medians and means, respectively. * indicates p < 0.05 after FDR correction for multiple comparisons.

### Functional significance of pattern persistence following Task B

Analogous to above, we assessed whether the hippocampal persistence following Task B reflected the allocentric or egocentric representations of the task. We therefore tested whether Task B/RS3 similarity differed between representations. Results showed that the similarity between RS3 and Task B was significantly greater for the allocentric as compared to the egocentric representation (t(52) =2.04, p(unc) = 0.047, Cohen’s d = 0.56; Figure 5). This result indicates that the hippocampal multivoxel patterns reactivated during post-test RS3 reflect the allocentric representation of the sequence task. [Note that the putamen did not exhibit pattern persistence following Task B (i.e., see Figure 4) and thus was excluded from the analyses assessing functional significance.]

**Figure 5:**
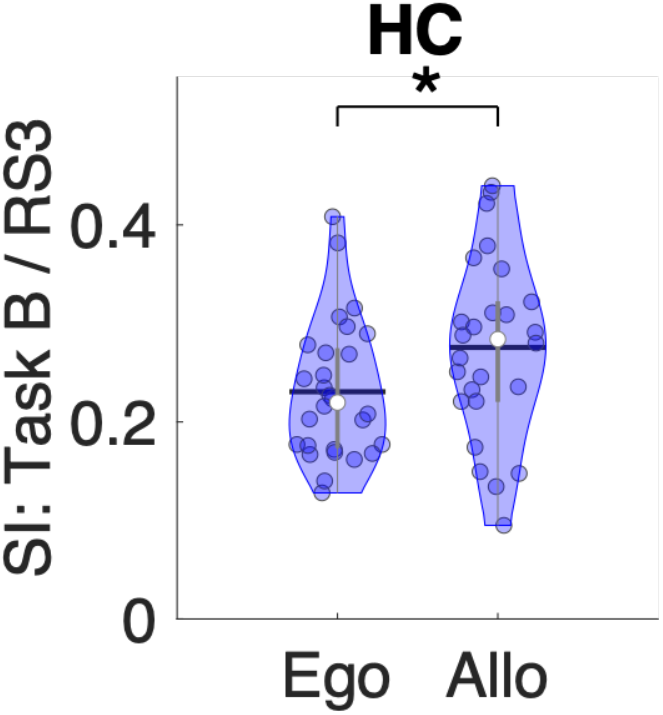
Functional significance of pattern persistence following Task B. Violin plots ^25^ depicting the Similarity Indices (SI) between MVCS matrices from the post-test RS (i.e., RS3) and those during Task B for the hippocampus (HC) separately for the egocentric (Ego) and allocentric (Allo) groups. SI was greater for the allocentric than the egocentric group, suggesting pattern persistence during RS3 (following Task B) reflects the allocentric representation of the motor sequence. For all violin plots, individual data points are shown as small colored circles jittered on the horizontal axis within the respective plot to increase visualization (N = 54; one subject was excluded due to missing data); white circles and horizontal black lines depict group medians and means, respectively. * indicates p < 0.05.

## Discussion

In this study, we used multivariate analyses of fMRI data acquired during a motor task and resting state to investigate whether brain patterns elicited by MSL persist into immediate post-learning waking rest. Results indicated that task-related patterns within the hippocampus and the striatum - but not in M1 - were reinstated during subsequent rest and thus provide evidence for the spontaneous reactivation of learning-related activity in awake rest epochs following motor sequence learning. Moreover, pattern persistence in the hippocampus specifically reflected the allocentric (spatial) representation of the acquired sequence, providing insights into the functional significance of hippocampal reactivation following motor learning.

Persistence of task-related hippocampal patterns into post-learning rest has been consistently observed in humans in the declarative memory domain ^1,4–6,27–30^. It is argued that such pattern persistence reflects “reactivation” or “replay” of patterns that were previously expressed during learning ^1^. The current study reports task-related hippocampal pattern persistence into post-learning rest in the motor memory domain. These findings extend previous neuroimaging investigations highlighting the crucial role of the hippocampus in motor learning [e.g., ^9,10^]. Moreover, our findings are in line with recent examinations assessing hippocampal activity on the micro-offline time scale (i.e., during rest blocks interleaved with practice). Jacobacci et al. ^17^ demonstrated that hippocampal activations during these micro-offline rest epochs - as revealed by univariate analyses - tended to be linked to the amplitude of micro-offline gains in performance as well as learning-related changes in hippocampal structure. Additionally, Buch and colleagues ^18^ conducted multivariate pattern analyses of magnetoencephalography (MEG) data and their results demonstrated hippocampal replay during micro-offline epochs. On the macro-offline timescale (i.e., between sessions of motor task practice), results from Buch et al. ^18^ indicated that the replay patterns in the post-task RS epoch were comparable to those observed pre-task and thus evidence of replay or pattern persistence into post-learning rest was lacking ^18^. The reasons for these discrepant findings between this earlier research and the current results are not immediately clear, but it is worth noting that the two studies employed different imaging modalities (MEG vs. fMRI) with inherently different spatial and temporal resolutions and distinct analytic approaches.

Earlier research from our own group in which a motor sequence task was manipulated to assess different task representations demonstrated that the hippocampus supports the allocentric (i.e., spatial) representation of acquired sequences ^26^. In line with this finding, results from the current study suggest that the spatial representation of an acquired motor sequence is specifically reactivated in the hippocampus during the early consolidation window following learning. These results are in line with a plethora of rodent studies suggesting that hippocampal replays consolidate previously learned spatial relationships (see ^19^ or a review). The current data thus provide novel insights into the functional significance of hippocampal reactivation following motor sequence learning in humans. That is, the offline reactivation of learning-related hippocampal activity reflects the spatial representation of the motor sequence acquired during online task practice.

The recurrence of task patterns into subsequent sleep and wakeful rest has been previously reported in rodent research in the ventral striatum [after reward-based learning; ^31^] and putamen [after exploration of novel objects; ^32^]. While there is evidence from previous human research that striatal multivariate patterns are modulated during motor learning ^16,33,34^, the current research reports persistence of motor task-specific striatal patterns into post-learning rest in humans. Interestingly, such reactivation or reverberation of striatal task patterns has been observed in rodents in conjunction with hippocampal replay ^21,32^. Even though there is no causal evidence that the hippocampus drives replay in the striatum, it has previously been argued that the hippocampus plays a network synchronization role by orchestrating memory trace consolidation in concert with other task-relevant areas such as the striatum ^35^. Although our results cannot directly speak to the hippocampus as an orchestrator of such inter-regional communications, our data do demonstrate that both the hippocampus and striatum are reactivated during wakefulness following initial learning and thus are consistent with the aforementioned rodent work ^21,32^.

Against our expectations that pattern persistence in the striatum would reflect egocentric (motoric) representations of the motor task^20^, the current results indicated that persistence of learning-related activity in the putamen did not differ between allocentric or egocentric task conditions. The functional significance of putamen replay after motor learning therefore remains unclear. Based on earlier observations that the striatal system is involved in learning predictive stimulus-response associations (e.g., finger-key mapping, ^36^), one might argue that pattern persistence in the striatum might reflect such task-specific associations during RS2 which, by definition, is not shared with tasks requiring different stimulus-response associations (i.e., task B). This interpretation remains speculative and warrants further investigation. Interestingly, and in contrast to the hippocampus, our data did not provide evidence of pattern persistence following Task B (i.e., allocentric / egocentric test) in the striatum. Specifically, multivoxel activity patterns during Task B were equally similar to the preceding and subsequent RS scans (i.e., RS2 and RS3). It is possible that this null result can be explained by the shorter amount of sequence practice in Task B (i.e., 6 blocks as compared to 14 blocks of Task A). Specifically, it could be argued that this practice duration was not sufficient to elicit the significant reinstatement of task-specific patterns of striatal activity in subsequent rest intervals. The current experimental design unfortunately does not allow us to test this potential explanation.

The results observed on M1 patterns stand in contrast with previous animal ^22^ and human ^23^ research showing replay of task-related M1 activity patterns during subsequent sleep. It could be argued that M1 reactivations preferentially occur during post-learning sleep as compared to wakefulness. However, this remains hypothetical, as earlier studies suggest that M1 does not play a prominent role in sleep-dependent motor memory consolidation [see ^8^ for a review]. Our results are in line with recent evidence showing that, in contrast to premotor areas, activity patterns in M1 do not show motor sequence representations but rather exhibit patterns related to the execution of individual finger movements ^37,38^. It is thus tempting to speculate that M1 patterns do not code for the motor sequence memory trace *per se*, which might explain the lack of M1 pattern reactivation during post-learning rest reported in the current study.

In contrast to previous research in the declarative domain ^4^, the current results cannot speak to whether the magnitude of the pattern persistence is related to longer-term memory consolidation or retention processes. It would thus be beneficial for future research to examine whether the reinstatement of task-relevant patterns following motor sequence learning is linked to long-term advantages at the behavioral level. Moreover, although the present data demonstrate pattern persistence during *wake* intervals following motor learning, it is unclear whether similar reactivation processes occur during post-learning *sleep* epochs. Given the known role of post-learning sleep in motor memory consolidation processes ^7^ as well as the original place cell work in animal models ^39,40^, one could expect to observe similar reactivation processes during sleep. It would be interesting for future research to examine this possibility directly.

In conclusion, our results show that patterns of brain activity within the hippocampus and striatum – but not M1 - observed during motor learning persisted into post-learning rest. Importantly, our data suggest that the hippocampus specifically replays the spatial representation of the learned motor sequence. Our results thus provide insights into the functional significance of brain reactivation after motor sequence learning in early consolidation windows consisting of wakefulness. Moreover, our findings add to the growing body of literature demonstrating that the hippocampus plays a vital role in learning and memory processes in the motor memory domain.

## Methods

### Participants

Sixty-two young, right-handed ^41^, healthy volunteers were recruited by local advertisements to participate in this study, which was approved by the Research ethics board of the “Regroupement en Neuroimagerie du Québec (RNQ), Centre de recherche de l’Institut universitaire de gériatrie de Montréal, Université de Montréal”. Participants did not suffer from any known psychological, psychiatric (including anxiety ^42^ and depression ^43^) or neurological disorders, and reported normal sleep during the month preceding the experiment, as assessed with the Pittsburgh Sleep Quality Index, ^44^. None of the subjects were taking medications at the time of testing. Moreover, none received formal training on a musical instrument or as a typist. All participants were asked to follow a 4-day constant sleep schedule (according to their own rhythm ± 1 h) before the experiment. Compliance to the schedule was assessed using both sleep diaries and wrist actigraphy measures (Actiwatch AW2, Bio-Lynx scientific equipment Inc., Montréal, Canada).

Of the 62 participants recruited for the study, seven were discarded from all analyses. Three participants were excluded for excessive head motion during functional brain imaging. Specifically, 2 participants had maximum linear translations that exceeded 3mm (~1 voxel) and an additional participant had more than 50% of the volumes removed as part of the scrubbing procedure (see details below). One participant was excluded because missing data from RS 1 and 2 scans (see Figure 1), precluding the assessment of persistence following initial learning. Three additional participants were excluded as they presented outlier similarity indices assessing Task A pattern persistence as part of the multivoxel imaging analyses described below (i.e., > 3 standard deviations away from the group average). Consequently, a total of 55 subjects were included (mean age ± SD: 23.8 ± 3.4 years, 36 females).

### Experimental design

Participants were trained to perform a MSL task (see below) in an MRI scanner at approximately 11:30 am. Resting state (RS) functional scans were acquired immediately prior (RS1) and after (RS2) initial MSL training (referred to as Task A; see Figure 1A) as well as following (RS3) a task variant assessing allocentric or egocentric motor sequence representations (referred to as Task B; see below for additional details). Note that these 5 functional imaging runs were the first runs of a larger experimental protocol completed by the same participants that examined the effects of sleep on offline motor memory consolidation processes (see ^20^ for full protocol).

### Motor sequence learning paradigm

The sequential finger tapping task used in this research was coded in Cogent2000 (http://www.vislab.ucl.ac.uk/cogent.php) and implemented in MATLAB (MathWorks Inc., Sherbom, MA). Participants were instructed to tap a five-element finger sequence on an MR-compatible keyboard with their non-dominant (left) hand as fast and as accurately as possible (Figure 1A). The sequence to perform was explicitly provided to the participants prior to training. Blocks of task practice were interleaved with 15-second rest intervals. Practice and rest were indicated by green and red fixation crosses, respectively, on the center of the screen. During the rest blocks, participants were instructed to keep their fingers still and look at the red cross. The number and timing of each specific key press within each practice block was recorded (60 key presses, ideally corresponding to 12 correct sequences). This procedure effectively controlled the number of movements executed in each block. Yet, the duration of the practice blocks progressively decreased with learning, as subjects became faster at executing the 60 key presses. Performance on the motor sequence learning task was assessed with a measure of movement speed (time to perform a correct sequence in s). As accuracy (i.e., number of correct sequences per block) in this specific experiment was quite high (> 11 correct sequences per block on average; maximum = 12) and stable across blocks of practice (see ^20^), it was not considered further in the current research.

For Task A, all participants performed 14 blocks of the sequence 4-1-3-2-4, where 1 through 4 represent the index through little fingers. For Task B (6 blocks), the keyboard and the subject’s hand were turned upside down (see Figure 1A) and participants were assigned to one of two experimental conditions that differed based on whether the allocentric (n=26) or egocentric (n=29) representation of the motor sequence was probed (see ^20^). The egocentric representation was assessed by having the participants complete the same sequence of finger presses (i.e., 4-1-3-2-4); however, with the keyboard/hand turned upside down, the spatial locations of the movement responses were now novel. Conversely, for the allocentric representation, the spatial representation of the sequence was preserved by having the participants complete the mirrored finger sequence (i.e., 1-4-2-3-1).

### fMRI data acquisition and processing

#### Acquisition

Functional MRI-series were acquired using a 3.0 T TIM TRIO scanner system (Siemens, Erlangen, Germany), equipped with a 32-channel head coil. Multislice T2*-weighted fMRI images were obtained, during both resting state and task practice, with a gradient echo-planar sequence using axial slice orientation (TR = 2650 ms, TE = 30 ms, FA = 90°, 43 transverse slices, 3 mm slice thickness, 10% inter-slice gap, FoV = 220 × 220 mm2, matrix size = 64 × 64 × 43, voxel size = 3.4 × 3.4 × 3 mm^3^). Slices covering the whole brain were acquired along the z-axis in an ascending direction. During each resting state scan (6 min 40 s), participants were instructed to keep their eyes open and fixate on a white cross in the middle of a black screen. They were also asked to remain still and “not to think about anything in particular”. A structural T1-weighted 3D MP-RAGE sequence (TR = 2300 ms, TE = 2.98 ms, TI = 900 ms, FA = 9°, 176 slices, FoV = 256 × 256 mm^2^, matrix size = 256 × 256 × 176, voxel size = 1 × 1 × 1 mm^3^) was also acquired in all subjects. Head movements were minimized using cushions.

#### Analyses

##### Pre-processing

The fMRI data were preprocessed using SPM12 (http://www.fil.ion.ucl.ac.uk/spm/software/spm12/; Wellcome Centre for Human Neuroimaging, London, UK). The reoriented structural image was segmented into gray matter (GM), white matter (WM), cerebrospinal fluid (CSF), bone, soft tissue, and background. Task-based and RS functional volumes of each participant were first slice-time corrected (reference: middle slice). Images were then realigned to the first image of each run and, in a second step, realigned to the mean functional image computed across all the individuals’ fMRI runs using rigid body transformations. The mean functional image was co-registered to the high-resolution T1-weighted anatomical image using a rigid body transformation optimized to maximize the normalized mutual information between the two images. The resulting co-registration parameters were then applied to the realigned functional images. To optimize voxel pattern analyses, functional and anatomical data remained in subject-specific (i.e., native) space, and no spatial smoothing was applied to functional images ^4^.

##### ROI definition

Three ROIs were considered in the analyses (see Figure 2B) and consisted of the bilateral hippocampus, bilateral putamen and right M1 (i.e., contralateral to the hand practicing the task). The hippocampus and putamen ROIs were created in the native space of each individual using the FMRIB’s Integrated Registration Segmentation Toolkit (FSL FIRST; http://fsl.fmrib.ox.ac.uk/fsl/fslwiki/FIRST). The bilateral hippocampal ROI included an average of 309 voxels (SD = 33; range of 258 – 385) and the bilateral striatal ROI included a mean of 309 voxels (SD = 22; range of 274 – 362). The right M1 ROI was defined by masking the activation map from the group-level main effect of practice map (task > rest) derived from univariate analyses (threshold p <0.05 whole brain FWE corrected) with the anatomically-defined right M1 from the Human Motor Area Template (HMAT) ^45^. Details on the univariate analytic pipeline and corresponding group-level results are provided in ^20^. As this functional map and anatomical template were provided in MNI space, the M1 ROI was mapped back to native space using the individual’s inverse deformation field output from segmentation of the anatomical image. The right M1 ROI included an average of 254 voxels (SD = 23; range of 204 – 302).

##### Multi-voxel correlation structure (MVCS) analyses

The custom analysis pipeline was written in Matlab. Prior to running MVCS analyses, additional preprocessing of the time series was completed. Specifically, whole-brain signal was detrended and high-pass filtered (cutoff=1/128). The realignment parameters were used to compute framewise displacement (FD). If the FD of any volume *n* exceeded 0.5mm, volume *n* and *n*+1 were discarded from analyses. The percentage of volumes excluded from the RS1, Task B, RS2, Task B and RS3 runs were 0.70 (SD=1.58), 5.90 (SD=8.06), 1.63 (SD=3.22), 7.68 (SD=9.54) and 1.33 (SD = 2.60), respectively. At the ROI level, only voxels with >10% GM probability were included in the analyses. Regression analyses were performed on the fMRI time-series of the remaining voxels in each ROI to remove additional sources of noise. These nuisance regressors included the first three principal components of the signal extracted from the WM and CSF masks created during segmentation of the anatomical image, the 6-dimensional head motion realignment parameters, as well as the realignment parameters squared, their derivatives, and the squared of the derivatives (i.e., a total of 30 nuisance regressors). Last, the number of volumes included in each specific analysis was matched across relevant runs within each individual. To do so, the run with the smallest number of volumes was identified, and for the other subset runs, the corresponding number of volumes centered around the middle volume of that run were selected. For the analysis assessing the persistence following Task A (results presented in Figure 2), the average number of volumes per run included in the analyses was 146 (SD = 6; range 117-150). The remaining analyses included Task B, which consisted of only 6 blocks of motor task practice and thus fewer number of volumes were obtained. For these analyses, the average number of volumes per run included was 94 (SD = 2; range 55-147).

Multi-voxel correlation structure (MVCS) matrices were computed for each ROI and each run of interest with similar procedures as in previous research ^4,6^. Specifically, Pearson’s correlations were computed between each of *n* BOLD-fMRI voxel time courses, yielding an *n* by *n* MVCS matrix per ROI, run and individual (see Figure 2A). Pearson’s correlation coefficients were Fisher z-transformed to ensure normality. Each MVCS matrix is thought to reflect the response pattern of the ROI in the specific run ^4^.

##### Statistical Analyses

To ensure participants learned the novel motor sequence, the behavioral measure of movement speed was assessed with one-way repeated-measures ANOVAs, with block as the within-subject factor for both Task A and Task B. As Task B consisted of a motor task variant that assessed either the allocentric or egocentric sequence representation, block x representation ANOVAs were conducted to ensure no performance differences between the two groups (as indicated in the initial analyses reported in ^20^). For all behavioral analyses, Greenhouse-Geisser corrections were applied if the sphericity assumption was violated.

To assess the influence of motor learning on multivoxel patterns within our ROIs, similarity indices (SI) that reflect the similarity of the multivoxel patterns between two specific fMRI runs were computed as the r-to-z transformed correlation between the two MVCS matrices of interest ^4,27^. Four statistical analyses were then conducted to test specific research questions of interest. *First*, to assess persistence of task-related patterns following Task A (i.e., initial motor sequence learning), SIs were computed between (a) RS1 and Task A and (b) Task A and RS2. The magnitude of the similarity indices between Task A and RS2 reflects the degree of persistence of the learning-related patterns into the subsequent rest interval. To assess whether this persistence is significant, a comparison was made to the SI between the Task A and RS1 (i.e., pre-training) as a within-subject control. Specifically, paired t-tests were computed using Matlab to compare SIs between RS1/Task A and Task A/RS2. Our *second* analysis assessed whether the observed pattern persistence revealed in the first analysis reflected the allocentric or egocentric representation of the motor sequence assessed by Task B. Specifically, independent t-tests were conducted to assess group differences (allocentric vs. egocentric) in the similarity between the multivoxel patterns observed during RS2 and Task B. *Third*, we examined whether multivoxel patterns during Task B showed evidence of persistence into the subsequent rest interval (i.e., RS3). This was achieved by conducting paired t-tests between Task B similarity with RS2 vs. RS3 across task variants. *Last*, and mirroring our second analysis described above, we assessed whether persistence following Task B (i.e., in RS3) reflected the allocentric or egocentric representation of the motor sequence. Accordingly, independent t-tests were conducted to assess group differences (allocentric vs. egocentric) in the similarity between the multivoxel patterns observed during RS3 and Task B.

Cohen’s d and partial η^2^ effect sizes corresponding to paired t-tests and ANOVAs, respectively, are reported. Within each family of hypothesis tests, results were considered significant if *p* < 0.05, with a false-discovery rate (FDR) correction for testing multiple ROIs ^46^.

## Data Availability

The approval granted by the local ethics committee does not permit the sharing of individual raw data. All source data underlying the figures shown in the main text are available as Supplementary Data.

## Code Availability

Scripts used for data analyses are available from the corresponding author upon reasonable request.

## Acknowledgements

This study was supported by grants from the Canadian Institutes of Health and Research (CIHR; MOP 97830) as well as the Ministry of Economic Development, Innovation and Exportation of Quebec (MDEIE; PSR-SIIRI-704) to JD. MAG received salary support from the Research Foundation Flanders (FWO; project G099516N and predoctoral fellowship 1141320N).

## Author Contributions

GA and JD conceived of and designed the experiment. GA conducted the experiments. GA, BRK, MAG and DM contributed to analytic tools. BRK analyzed the data. BRK and GA wrote the initial draft of the manuscript. All authors contributed to the subsequent revisions.

